# SpeciateIT and vSpeciateDB: Novel, fast and accurate per sequence 16S rRNA gene taxonomic classification of vaginal microbiota

**DOI:** 10.1101/2024.04.18.590089

**Authors:** Johanna B. Holm, Pawel Gajer, Jacques Ravel

## Abstract

Clustering of sequences into operational taxonomic units (OTUs) and denoising methods are a mainstream stopgap to taxonomically classifying large numbers of 16S rRNA gene sequences. We developed speciateIT, a novel taxonomic classification tool which rapidly and accurately classifies individual amplicon sequences (https://github.com/Ravel-Laboratory/speciateIT). Environment-specific reference databases generally yield optimal taxonomic assignment. To this end, we also present vSpeciateDB, a custom reference database for the taxonomic classification of 16S rRNA gene amplicon sequences from vaginal microbiota. We show that speciateIT requires minimal computational resources relative to other algorithms and, when combined with vSpeciateDB, affords accurate species level classification in an environment-specific manner.

**Importance:** Herein, two resources with new and practical importance are described. The novel classification algorithm, speciateIT, is based on 7^th^ order Markov chain models and allows for fast and accurate per-sequence taxonomic assignments (as little as 10 min for 10^7^ sequences). vSpeciateDB, a meticulously tailored reference database, stands as a vital and pragmatic contribution. Its significance lies in the superiority of this environment-specific database to provide more species-resolution over its universal counterparts.

## Introduction

High-throughput next generation sequencing has revolutionized the field of metataxonomics by producing millions of sequences at an affordable cost, increasing the depth at which microbial communities are characterized. However, large sequence datasets have led to new challenges such as high computational costs associated with data analyses and accurate taxonomic classification. Bioinformaticists have developed novel sequence clustering algorithms which either produce, for a given similarity threshold, groups of sequences known as operational taxonomic units (OTUs) [2-4], or reduce sequencing errors and minimize noise in the data [5-8]. These approaches have proven useful, though noise reduction/error correction may artificially remove or produce diversity [7] and the process of OTU clustering simply shifts the computational cost from taxonomic classification assignment to clustering and is not without problems. Most significantly, the transitive taxonomic assignments obtained from an OTU representative sequence are often flawed as 10-30% of sequences within the OTU, if processed separately, are assigned different taxonomy, thus challenging the ecological value of an OTU [9]. Further, output from clustering-based analyses are dataset specific and when data are added to a study, clustering must be run again and at increasing computational cost.

To alleviate these issues, we developed speciateIT, an algorithm capable of fast, accurate individual sequence taxonomic classification. Using a model guide tree and 7^th^ order Markov chain models to represent bacterial species trained on taxonomy-adjusted amplicon specific regions sequences, speciateIT requires little computational resources, and can quickly process large sequence datasets.

Additionally, environment-specific reference databases improve species-level classification accuracy and precision by reducing misclassification to species irrelevant to the environment and increasing study reproducibility and generalizability to other studies [10, 11]. For these reasons, we have also developed vSpeciateDB, a set of custom databases of reference sequences for classifying vaginal microbiota. SpeciateIT models for the vaginal microbiota correctly classified to the species level 99.9% of training set 16S rRNA gene V1-V3, V3-V4, and V4 regions sequences. This is a major improvement over the RDP Naïve Bayesian Classifier [12], which is capable of 99% classification accuracy of known sequences and 76% of novel sequences (not part of the training data) to the genus-level [13, 14].

## Results

Estimates of classification accuracy for novel sequences were obtained using 10-fold cross validation. To ensure confidence in assignments, speciateIT imposes model-specific classification error thresholds: when the posterior probability of a query sequence does not exceed this threshold, the query sequence is classified as the next highest taxonomic level at which this threshold requirement is met. In the case of novel taxa, speciateIT is expected to assign higher-level classifications. In 10-fold cross-validation testing, 98.7, 97.6, and 97.2% of sequences from “known” species (a species with at least 1 sequence present in the training dataset) were correctly assigned with >90% of assignments made to the genus or species levels (**Figure 1A**). For sequences from “novel” species (those with no sequences present in the training set), 60-70% were correctly assigned to their respective taxonomic categories, with the accuracy varying depending on the targeted region. This highlights the efficacy of speciateIT in accurately classifying bacterial taxa using higher order Markov chain models.

**Figure 1.**
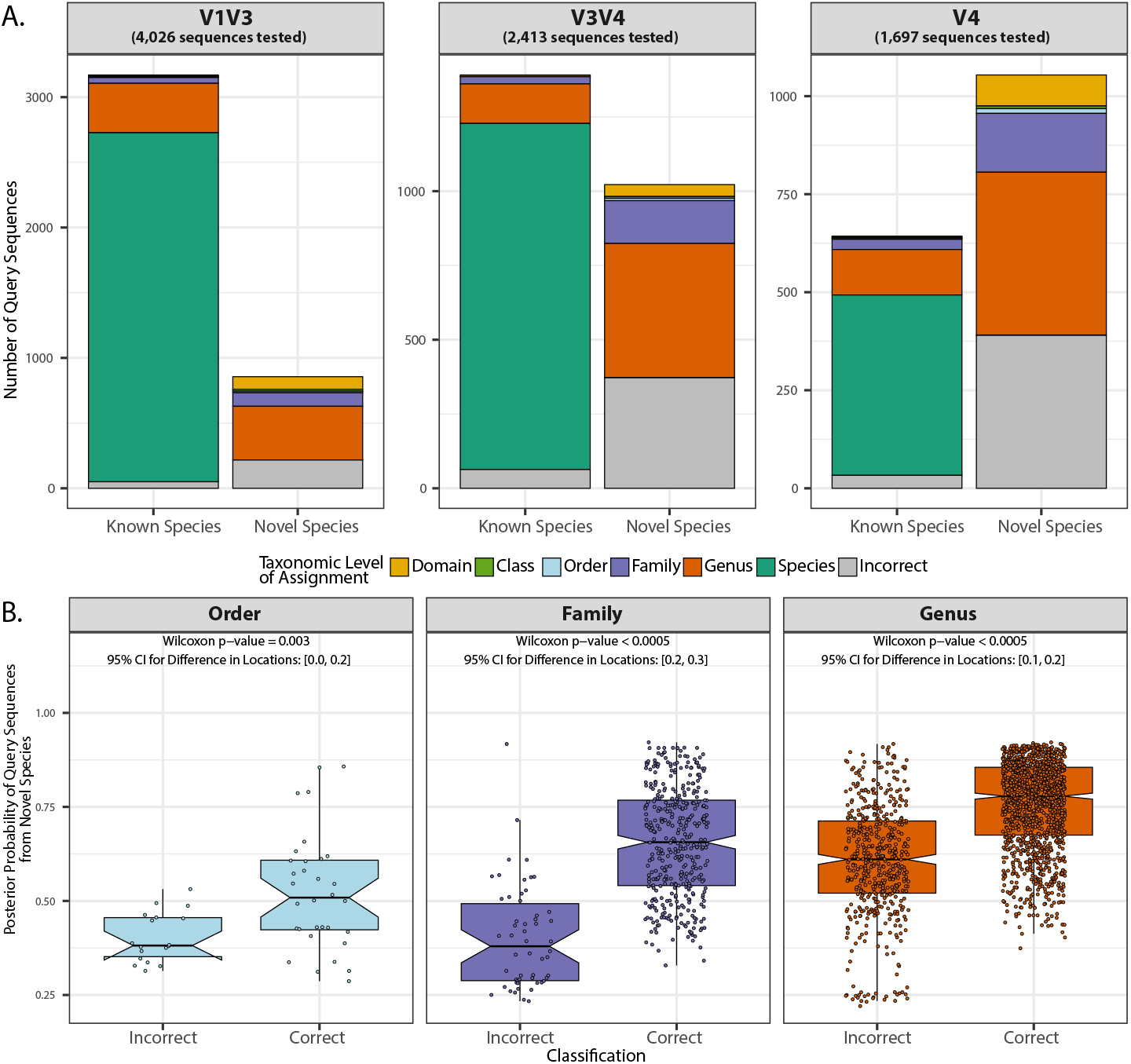
(A) Ten-fold cross validation of the vSpeciateDB V1V3, V3V4, and V4 models demonstrated exceptional classification of sequences from “Known Species” with at least 1 sequence present in models. Most sequences from “Novel Species” were correctly classified at some taxonomic level. (B) The posterior probabilities of query sequences from “Novel Species” tended to be higher for correct classifications relative to incorrect classifications.

An essential aspect of speciateIT is its provision of posterior probabilities for query sequences. In the context of classification using Markov chain models, posterior probabilities represent the likelihood or confidence that a given sequence belongs to a particular category or class. These probabilities are calculated based on the observed sequence data and the parameters of the Markov chain model. When a query sequence has a lower posterior probability, it suggests that the observed sequence data is less consistent with the model’s parameters. This can indicate that the sequence deviates more from the typical patterns captured by the model, potentially suggesting a poorer match between the sequence and the model. However, it’s important to note that a lower posterior probability does not necessarily mean that the classification result is incorrect or that the sequence is not related to the modeled categories. It simply suggests lower confidence in the classification result. In some cases, a sequence with a lower posterior probability may still be correctly classified, especially if the model captures only part of the variability present in the data. Regarding sequences from novel species (those absent from the training set), cross-validation results illustrate that the posterior probabilities from correct genus-, family-, or order-level assignments tend to be greater than incorrect classifications (**Figure 1B**).

To compare classification of vaginal microbiota using SpeciateIT with vagina-specific vSpeciateDB to other popular classifiers and reference sets (RDP Naïve Bayesian Classifier stand-alone Bioconda version 2.13, default settings; DADA2 implementation of RDP Naïve Bayesian Classifier trained with SILVA v138.1 and GTDB r86 reference sets), we classified independent sequences from GTDB (not included in the production of vSpeciateDB) truncated to each variable region and included those from the 100 most abundant species detected in the vaginal microbiota [1]. SpeciateIT with vSpeciateDB provided more species-level assignments than other methods including the DADA2 implementation of the RDP classifier which provided species level assignments, when possible (function: addSpecies) (**Figure 2**).

**Figure 2.**
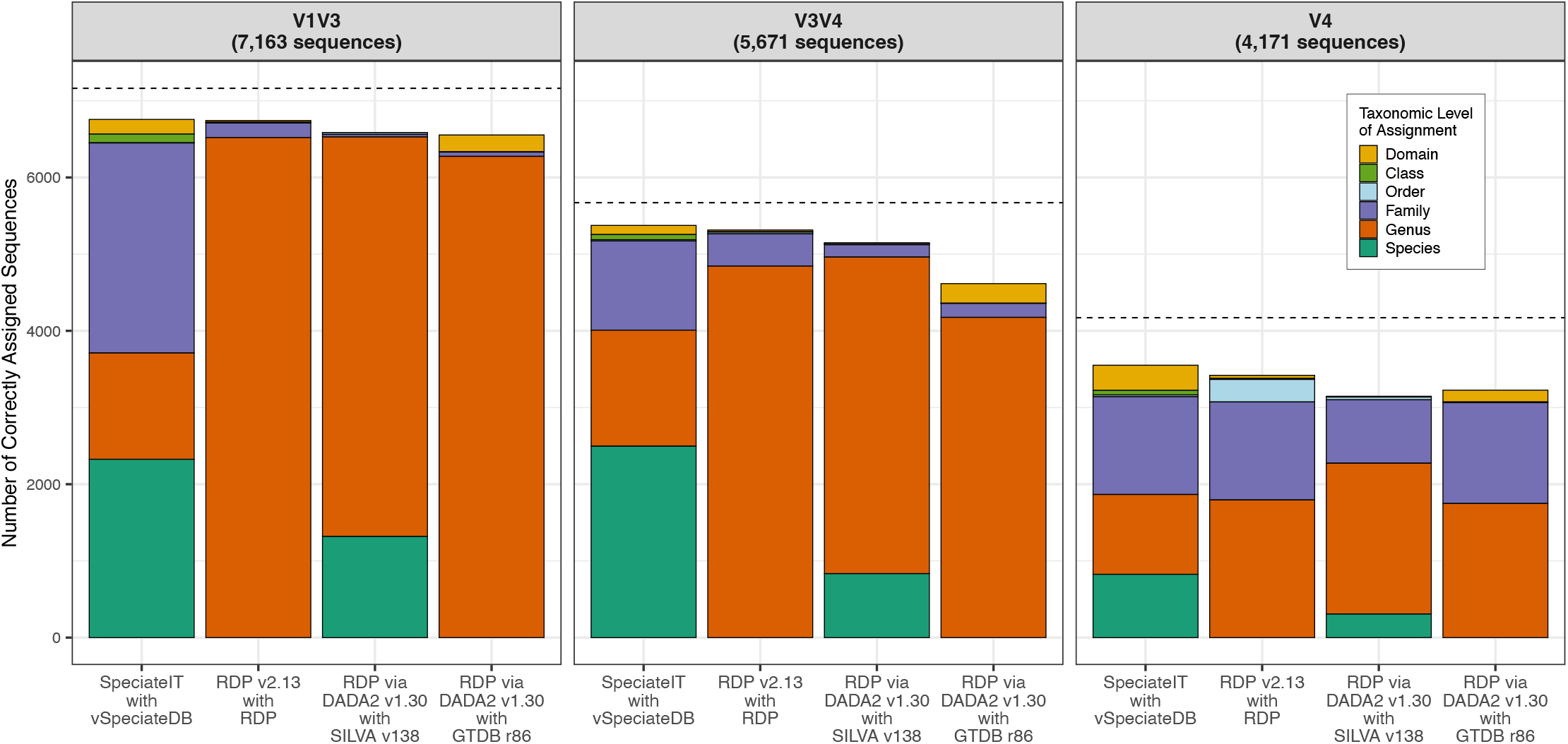
SpeciateIT outperforms other classification methods by providing correct species-level assignments. Dashed lines indicate total number of sequences tested.

The speed of speciateIT is incomparable because of its novel model tree-based approach which directs query sequence classification from the top of the tree (Root) to the branch or node of its final classification (**Figure S1**). Classification speed was measured on a 2021 Macbook Pro with an Apple M1 Max processor and 64G RAM using each amplicon reference training set sampled to 10^1^-10^7^ sequences and processed on one core. We compared the speed of speciateIT classification to the RDP Naïve Bayesian Classifier (stand-alone Bioconda version 2.13, default settings). SpeciateIT classified 1 million sequences in 3, 2, and 1 min for the V1V3, V3V4, and V4 classifiers, respectively (**Figure 3**). Speed is dependent on the number of models read for each classifier (the V4 classifier represents fewer species and therefore contains fewer models). Comparatively, the RDP Classifier classified 1 million V1-V3, V3-V4, and V4 sequences in 66, 57, and 32 min, respectively.

**Figure 3.**
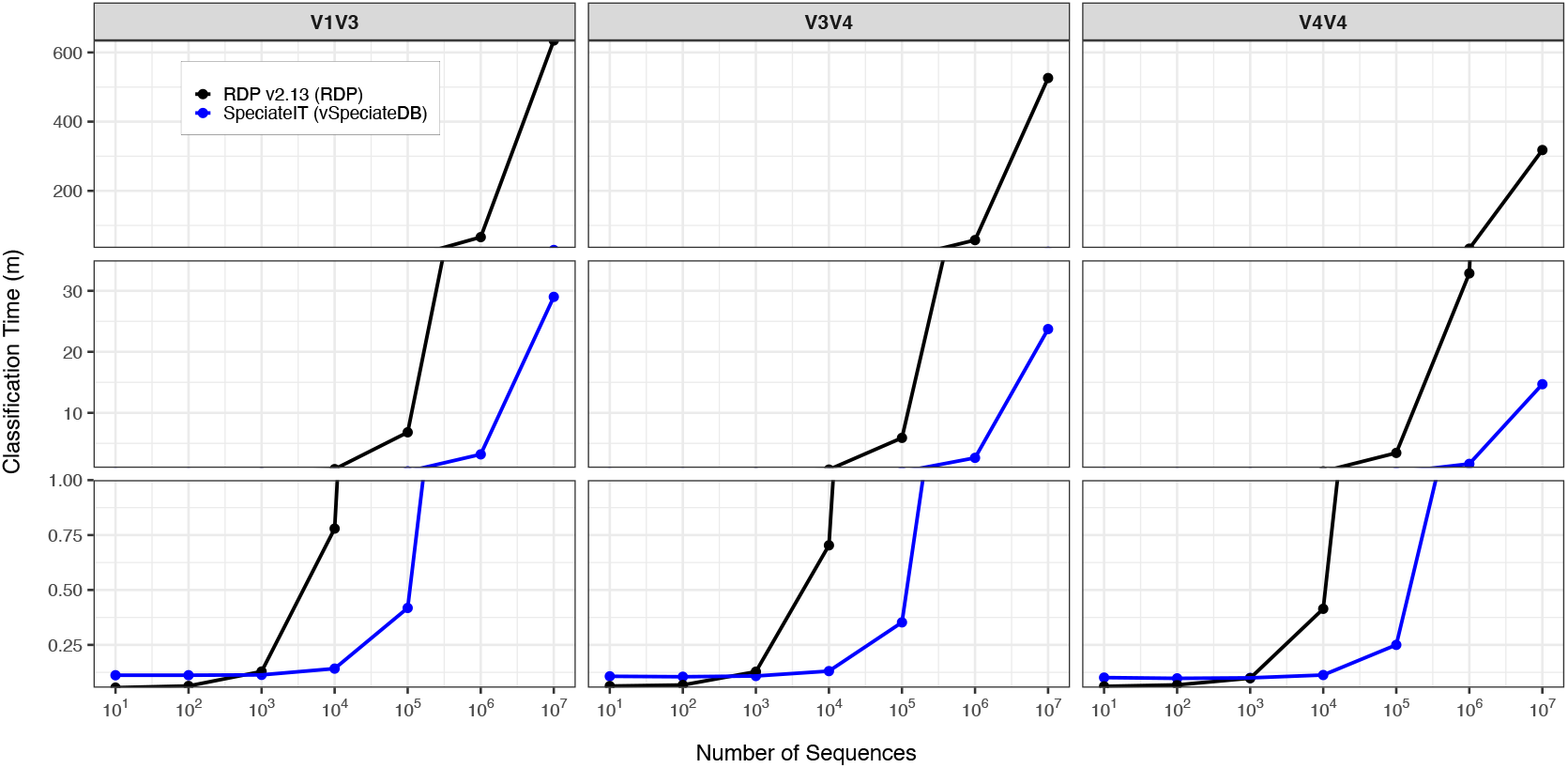
SpeciateIT is faster than the RDP classifier when datasets are greater than 1,000 sequences.

The performance of any classifier is entirely dependent on the quality of the sequence training set used to build it. Currently, speciateIT models have been built from full length 16S rRNA gene sequences curated from the Genome Taxonomy Database (GTDB) for the taxonomy-adjusted V1-V3, V3-V4, and V4 amplicon sequence regions for vaginal microbiota, and are publically available (https://github.com/ravel-lab/speciateIT). The full-length database comprises 2,224 species, 497 genera, 77 families, 36 orders, 16 classes, and 14 phyla.

One recent change in the field of vaginal microbiota is the expansion of *Gardnerella vaginalis* to multiple species. Eleven species are represented in the GTDB SSU rRNA reference sequence set from which vSpeciateDB sequences originated. *G. vaginalis* C was not included in the final training sets because no reference sequences contained the V3 or V4 regions. *Gardnerella vaginalis A* and *Gardnerella vaginalis F* were distinct from other *Gardnerella* species in both the V2 and V4 regions (**Figure S2a**). It was not possible to confidently distinguish other *Gardnerella* species at any targeted region. To maintain simplicity, one *Gardnerella* model (“*G. vaginalis”*) represents GTDB species: *G. leopoldii, G. piotii, G. swidsinskii, G. vaginalis and G. vaginalis* A, B, C, D, E, F, and H combined. Of other prevalent species in the vaginal microbiota, *Lactobacillus iners, L. jensenii, L. mulieris*, and “*Ca*. Lachnocurva vaginae” were distinct in vSpeciateDB while *L. gasseri* and *L. paragasseri* were not distinguishable at any region and are referred to as only *L. gasseri*. Notably, *L. crispatus* and *L. acidophilus* were indistinguishable at the V4 region (**Figure S2b**). Because *L. crispatus* is arguably more prevalent in the vaginal microbiota, these models are referred to as *L. crispatus*.

Lastly, the VAginaL community state typE Nearest CentroId classifier (VALENCIA) uses reference centroids representing microbiota compositions for each CST. The taxonomic annotations used in building the reference centroids are integral to correct CST classification. Because vSpeciateDB-based taxonomic assignments differ from those used in the current version of VALENCIA reference centroids, we have produced reference centroids based on vSpeciateDB taxonomy and compatible with the VALENCIA algorithm for CST assignment (**Figure S3**).

## Discussion

We anticipate vSpeciateDB will grow as more vaginal species are characterized, and more 16S rRNA gene variable region vSpeciateDB models will be produced. Importantly, taxonomy is adjusted in each V1-V3, V3-V4 and V4 database to reflect the loss of taxonomic information associated with sequence truncation, a known problem when using amplicon sequences [15]. Furthermore, the steps for vSpeciateDB curation are provided and can be used as a foundation upon which other environment-specific reference databases can be curated. The classification algorithm script, reference models, and VALENCIA reference version 2 centroids are available via https://github.com/Ravel-Laboratory/speciateIT.

## Materials and Methods

### speciateIT Algorithm

The core model building algorithm in speciateIT produces higher order Markov chain models for groups of phylogenetically related sequences. These groups were organized in a model tree reflecting the species lineages (buildModelTree). For each node of the model tree (except the root) a fasta file of all reference sequences corresponding to the node’s subtree was created and used to build Markov chain models (buildMC). Node-specific classification error thresholds were estimated to produce confidence in taxonomic assignments. During training set evaluation, if all sequences in a species were correctly assigned to the species, error thresholds were defined as the log10 of the minimum posterior probability minus 0.05 to allow flexibility in novel sequence assignment. If any sequences in a species were incorrectly assigned during training set evaluation, the error thresholds were defined as the log10 of the minimum posterior probability of correctly assigned sequences without added flexibility (err_thlds.R). Classification of a query sequence begins at the top of the model tree. The model producing the highest posterior probability is chosen, and the assignment is given to the sequence given that the posterior probability is greater than the classification error threshold for that model. Model comparisons at the next and lower taxonomic levels commence until either a terminal node (species-level) classification is reached, or the classification error threshold criterion is not met. All code presented here is available at https://github.com/ravel-lab/speciateIT and is included in speciateITv2.Rmd.

### speciateDB Curation

#### Building Reference Sequence Database

Environment-specific reference databases increase classification accuracy and study reproducibility [10, 11]. All code for reference database curation can be found in speciateITv2.Rmd. To build the vagina-specific vSpeciateDB reference database, we extracted sequences from the GTDB [16] small subunit rRNA gene sequence dataset (https://data.gtdb.ecogenomic.org/releases/release214/214.1/genomic_files_all/ssu_all_r214.tar.gz, referred herein as GTDB-SSU v214.1) for species found in the vaginal microbiome[17, 18] as per VIRGO2 (virgo.igs.umaryland.edu, and species list available on https://github.com/ravel-lab/speciateIT) resulting in 308,611 16S rRNA gene sequences from 14 bacterial phyla including 16 classes, 36 orders, 77 families, 497 genera, and 2,224 species (**Table 1**). Sequences were truncated to the V1-V4 region using tagcleaner.pl [19] with the 27F and 806R primer sequences allowing for 9 and 17 mismatches, respectively. Using mothur v.1.48.0 [20], truncated sequences were dereplicated after filtering those with ambiguous bases and lengths < 250 bp or > 1000 bp. RDP-formatted lineages were re-formatted (reversed and tab-delimited), and a taxonomy file was created connecting sequence IDs to species annotation.

**Table 1.** Summary of sequence information comprising each speciateIT 16S rRNA gene sequence region-specific database.

|  | Vaginal subset of GTDB-SSU v214.1 | V1-V3 | V3-V4 | V4 |
| --- | --- | --- | --- | --- |
| **Sequences** | 308,611 | 4,502 | 2,584 | 1,735 |
| **Species** | 2,224 | 1,322 | 1,272 | 1,165 |
| **Genera** | 497 | 406 | 415 | 411 |
| **Families** | 77 | 74 | 73 | 69 |
| **Orders** | 36 | 35 | 35 | 33 |
| **Classes** | 16 | 15 | 15 | 14 |
| **Phyla** | 14 | 13 | 13 | 12 |

#### Production of 16S rRNA gene sequence region-specific datasets

The final de-replicated V1-V4 dataset was used to produce datasets for the 16S rRNA gene amplicon V1-V3, V3-V4, and V4 regions using tagcleaner.pl (version 0.16) [19] and the 319F, 515F and 534R primers allowing for 9, 3, and 5 mismatches, respectively. V1-V3 and V3-V4 sequences were screened to remove sequences < 400 and > 500 bp long. V4 sequences were required to be 240-260 bp. Each dataset was then dereplicated using the unique.seqs command from mothur v.1.48.0 [20]. SpeciateIT models were constructed for each dataset and training set evaluation identified incorrectly classified sequences, which were subsequently removed (**Figure S4**). Most incorrect classifications at this stage originated from species over-represented in the reference database including *Escherichia coli, Klebsiella pneumoniae, Staphylococcus aureus, Salmonella enterica, Acinetobacter baumannii*, and *Streptococcus pneumoniae, S. epidermidis*, and *S. agalactiae*. Therefore, for these species and others with more than 50 sequences, three sequences with the highest posterior probabilities from training set evaluations were maintained. To further ensure quality of training data, sequence from each species aligned pairwise with Biostrings v2.70.2 [21], and sequences with less than 90% identity to all other sequences in the species were removed. Next, multiple sequence alignments for each database were produced with MAFFT v7.394 [22], and used to build phylograms with FastTree 2.1.10 [23]. Semi-supervised clustering using VI-cut [24] was performed and VI-cut clusters were evaluated for species purity and species indistinguishable by the targeted variable region(s). Within a region, if a cluster contained multiple species annotations, these annotations were merged and captured in the region’s concatenation map or “cat map”. All species annotations for models were replaced with the first alphanumeric species in the concatenation. When a species’ annotation was present in multiple nearby clusters (difference in cluster numbers ≤ 2), all species’ annotations within and between the clusters were concatenated. When species’ annotations were present in distant clusters, sequences in the smaller clusters were removed. The resulting vagina-specific datasets are collectively referred to as vSpeciateDB.

#### Testing classification accuracy of known and novel sequences

To estimate the capability of models to classify novel sequences, ten-fold cross-validations were performed on each vSpeciateDB (pecan_cv5.pl, available on GitHub). All curated sequences for a targeted region were included in the training set. Each dataset was randomly spit into training (90% of sequences) and test (10%) sets. SpeciateIT models were built from the training set and training set evaluation were performed for construction of error thresholds. Subsequently, the test set was classified.

#### Comparing classifications for multiple classifiers

Sequences from the GTDB-SSU v214.1 datasets that were truncated to the V1-V3, V3-V4, and V4 regions and excluded from vSpeciateDB were used as independent query sets to compare classification between (1) speciateIT trained with vSpeciateDB, (2) RDP Naïve Bayesian Classifier stand-alone Bioconda version 2.13 (default settings), and the DADA2 implementation of RDP Naïve Bayesian Classifier trained with (3) SILVA v138.1 or (4) GTDB r86 reference sets. For the DADA2 implementations, both the assignTaxonomy and addSpecies functions were employed. Classifications were compared to the taxonomy of the GTDB-SSU v214.1 dataset to determine correctness.

#### Speed of Classification

Random test sets of 10^1^-10^7^ sequences were produced from the reference sequences used to build models. Classification for each set was performed on 2021 Macbook Pro with an Apple M1 Max processor using 1 core. For RDP classification, the RDP Naïve Bayesian Classifier (stand-alone Bioconda version 2.13, default settings) was used (rdp_classifier classify -f allrank). For speciateIT classification, models and error thresholds for variable region-specific 16S rRNA gene sequences being classified were employed. Time of classification was measured using the time bash utility.

## Acknowledgments

Research reported in this publication was supported by the National Institute of Allergy and Infectious Diseases of the National Institutes of Health under award numbers F32-AI136400 (JH) and K01-AI163413 (JH), U19-AI158930 (JR), U19-AI084044 (JR), and R01-AI116799 (JR, Co-I).

## SUPPLEMENTAL FIGURES

**Figure S1.**
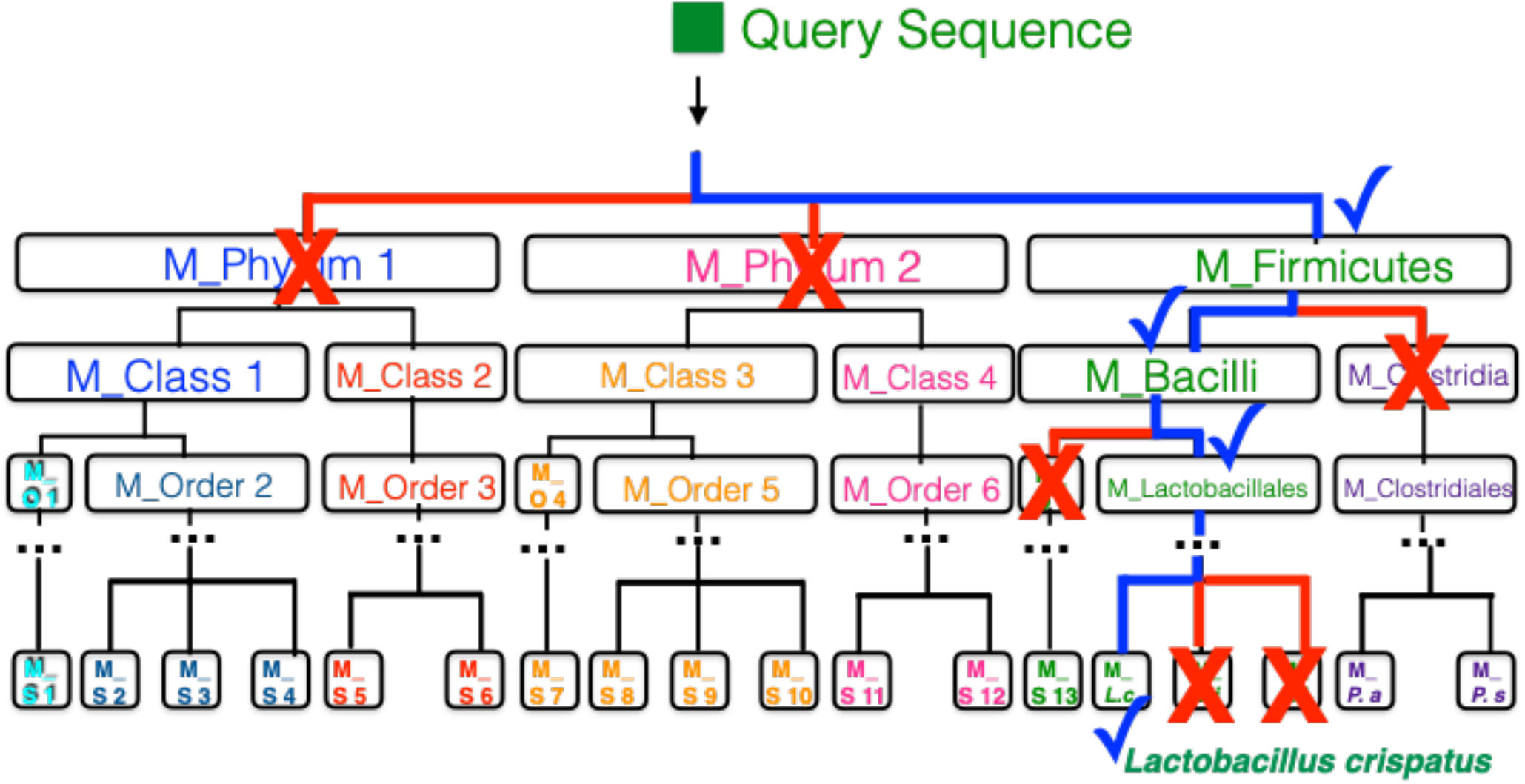
A taxonomy is assigned to a query sequence by traversing a Markov chain model tree associated with the taxonomic tree. First, a phylum level taxonomy is assigned, based on the posterior probability of the query sequence coming from the given phylum model. The process continues until a terminal node is reached or the error threshold requirement is not met. In this example, the query sequence is classified as *Lactobacillus crispatus* (*L*.*c*).

**Figure S2.**
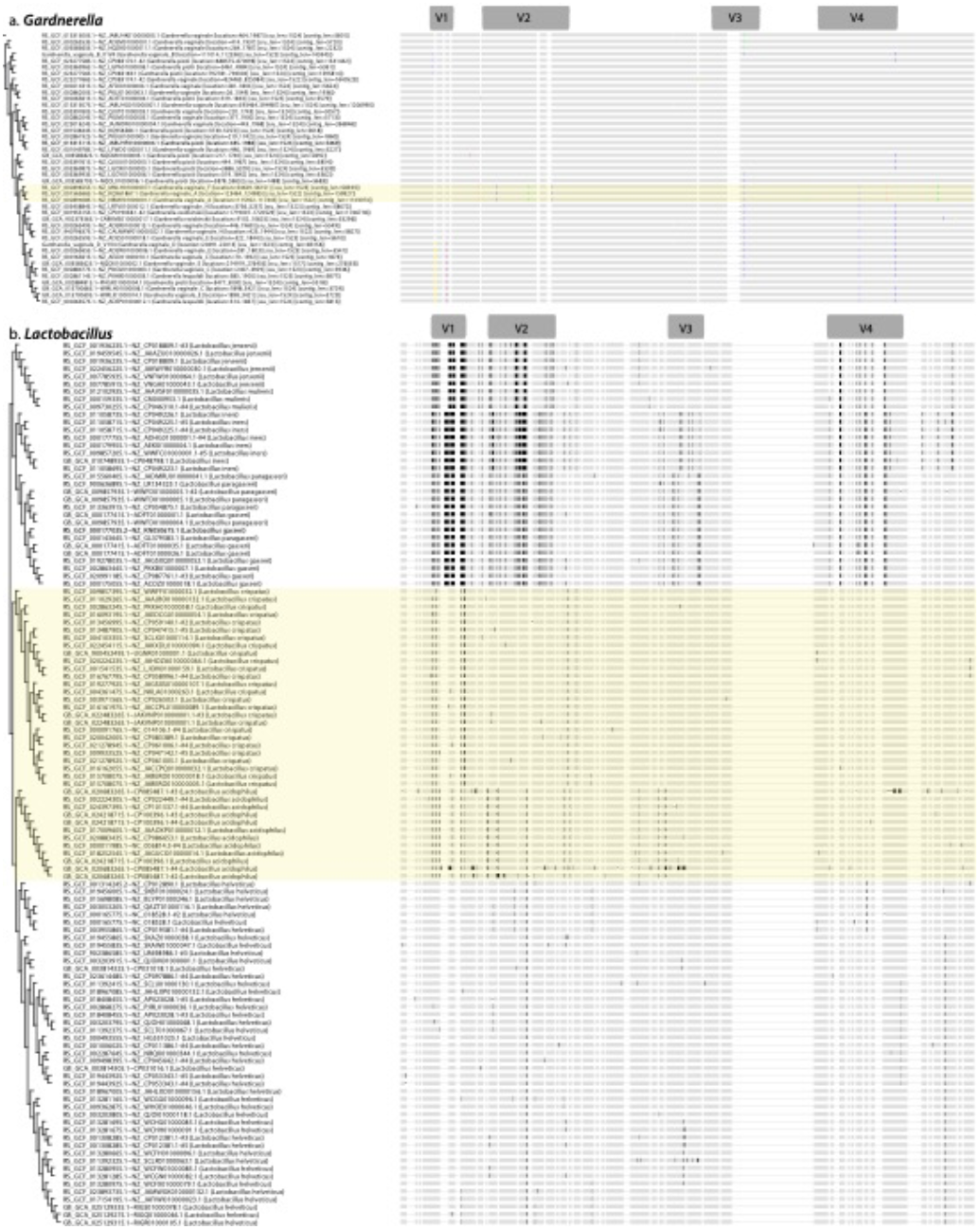
The V1-V4 regions of a) *Gardnerella* and b) *Lactobacillus* species 16S rRNA genes. *Gardnerella vaginalis* A and *G. vaginalis* F are distinct from other species at the V2 and V4 regions (highlighted in yellow). Other species are not distinguishable. b) *Lactobacillus* species are distinct except for *L. crispatus* and *L. acidophilus* at the V4 region.

**Figure S3.**
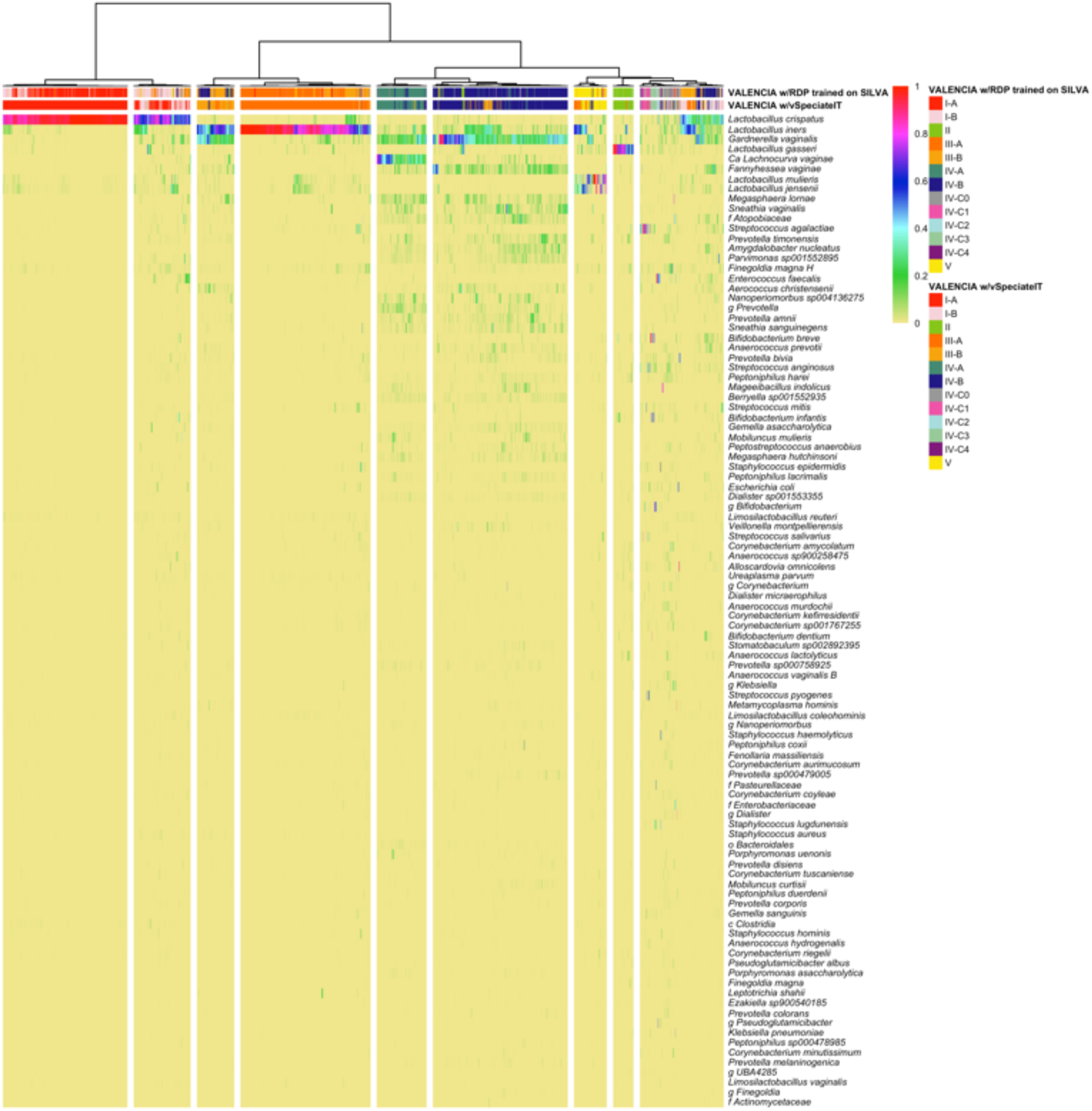
CST assignments by VALENCIA [1] using original reference centroids based on taxonomy from RDP classification trained on SILVA compared to the CST assignments by VALENCIA using centroids built from taxonomic assignments from speciateIT models trained on vSpeciateITDB (V3V4).

**Figure S4.**
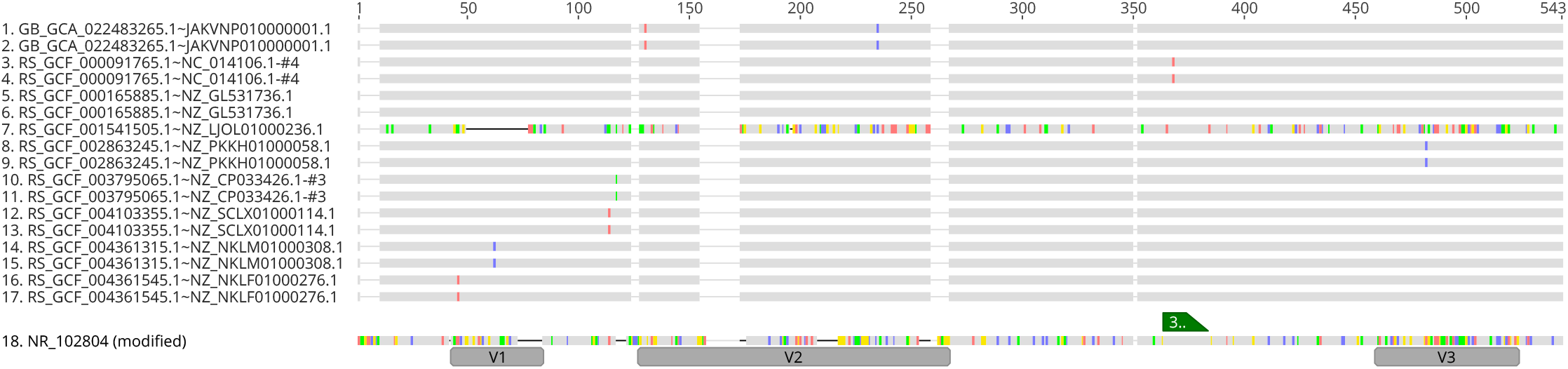
Incorrect classification during training set evaluation was an efficient, alignment-free method to cull sequences during training set curation. For example, sequence number 7 (RS_GCF_001541505.1∼NZ_LJOL01000236.1) was present in the training set following dereplication of sequences truncated to the V1-V3 region. During training set evaluation, RS_GCF_001541505.1∼NZ_LJOL01000236.1 was incorrectly classified to *Metamycoplasma hominis* instead of the expected assignment of *Lactobacillus crispatus*. The sequence was removed from all downstream analysis.

